# BMSS2: a unified database-driven modelling tool for systematic model selection and identifiability analysis

**DOI:** 10.1101/2021.02.23.432592

**Authors:** Russell Kai Jie Ngo, Jing Wui Yeoh, Gerald Horng Wei Fan, Wilbert Keat Siang Loh, Chueh Loo Poh

## Abstract

**Summary:** Modelling in Synthetic Biology constitutes a powerful tool in our continuous search for improved performance with rational Design-Build-Test-Learn approach. In particular, kinetic models unravel system dynamics, enabling system analysis for guiding experimental designs. However, a systematic yet modular pipeline that allows one to identify the “right” model and guide the experimental designs while tracing the entire model development and analysis is still lacking. Here, we introduce a unified python package, BMSS2, which offers the principal tools in model development and analysis—simulation, Bayesian parameter inference, global sensitivity analysis, with an emphasis on model selection, and a priori and a posteriori identifiability analysis. The whole package is database-driven to support interactive retrieving and storing of models for reusability. This allows ease of manipulation and deposition of models for the model selection and analysis process, thus enabling better utilization of models in guiding experimental designs.

**Availability and Implementation:** The python package and examples are available on https://github.com/EngBioNUS/BMSS2. A web page which allows users to browse and download the models (SBML format) in MBase is also available with the link provided on GitHub.

**Supplementary Information:** Supplementary data is available.

## Introduction

Modelling in synthetic biology guides experimental designs and enables rational optimization of the performance of biological systems (Elowitz, 2000; Jayaraman et al., 2016). In particular, kinetic modelling offers quantitative insights into the dynamic aspects and interactions of the systems components (Jayaraman et al., 2018). This has empowered researchers to improve the designs of genetic devices including biosensors (Holowko et al., 2016), cell factories (Yeoh et al., 2020), therapeutics (Caliendo et al., 2019), and integral feedback systems (Aoki et al., 2019).

Several useful tools supported by graphical user interface, e.g., iBioSim (Watanabe et al., 2019), COPASI (Hoops et al., 2006), and JWS Online (Olivier and Snoep, 2004), have been developed to facilitate the modelling process. In general, these tools provide key modelling features such as model building, simulation, sensitivity analysis, documentation in standard file formats and web-based storage. However, the model selection and identifiability analysis which are the prerequisites to shortlist the ‘right’ model and guide the experimental design are often not well undertaken, and yet it can be challenging to incorporate them readily into the existing tools. The community has geared toward the use of more programmable python-based packages which offer higher flexibility and support extensive scientific computation. These include SloppyCell (Myers et al., 2007) which analyses parameter uncertainties of model fitting, BMSS (Yeoh et al., 2019) which supports automated biomodel selection, and Tellurium (Choi et al., 2018) which covers simulation and standard files generation. Nevertheless, most of these efforts are focused on designated functionalities and thus limited in their scope.

Here we introduce BioModel Selection System (BMSS2), a refactored and extended version of BMSS (Yeoh et al., 2019), to provide a unified python library for model building, selection and analysis. The python package is designed to be modular, highly extensible, and open source to benefit from the active development of the scientific computing ecosystem and enable new functions to be readily integrated. Importantly, the whole package is database-driven to allow one to interactively retrieve and store models from/into SQL databases to ensure reproducibility and reusability, particularly for ease of model selection. In addition to supporting standard model creation and simulation, BMSS2 offers important features including parameter estimation using Bayesian inference (BI) based on Markov chain Monte Carlo (MCMC), trace analysis for *a posteriori* identifiability study, automated model selection based on Akaike Information Criteria (AIC), a priori structural identifiability using the STRIKE-GOLDD algorithm (Villaverde et al., 2019), and global sensitivity analysis for parameter tuning. To ensure interoperability, models can be exported in standard SBML formats.

## Design and Implementation

Our goal is to offer a unified platform to streamline the model building, selection, analysis, and interactive management and storage of kinetic models for biological systems and genetic devices via a database-driven approach (Figure 1). This is enabled by the unique feature of having both a global database (MBase) that provides a library of pre-built models and a local database (UBase) where users can easily manipulate and update their own models for model selection and analysis, encouraging the reuse and sharing of models among the community.

**Figure 1:**
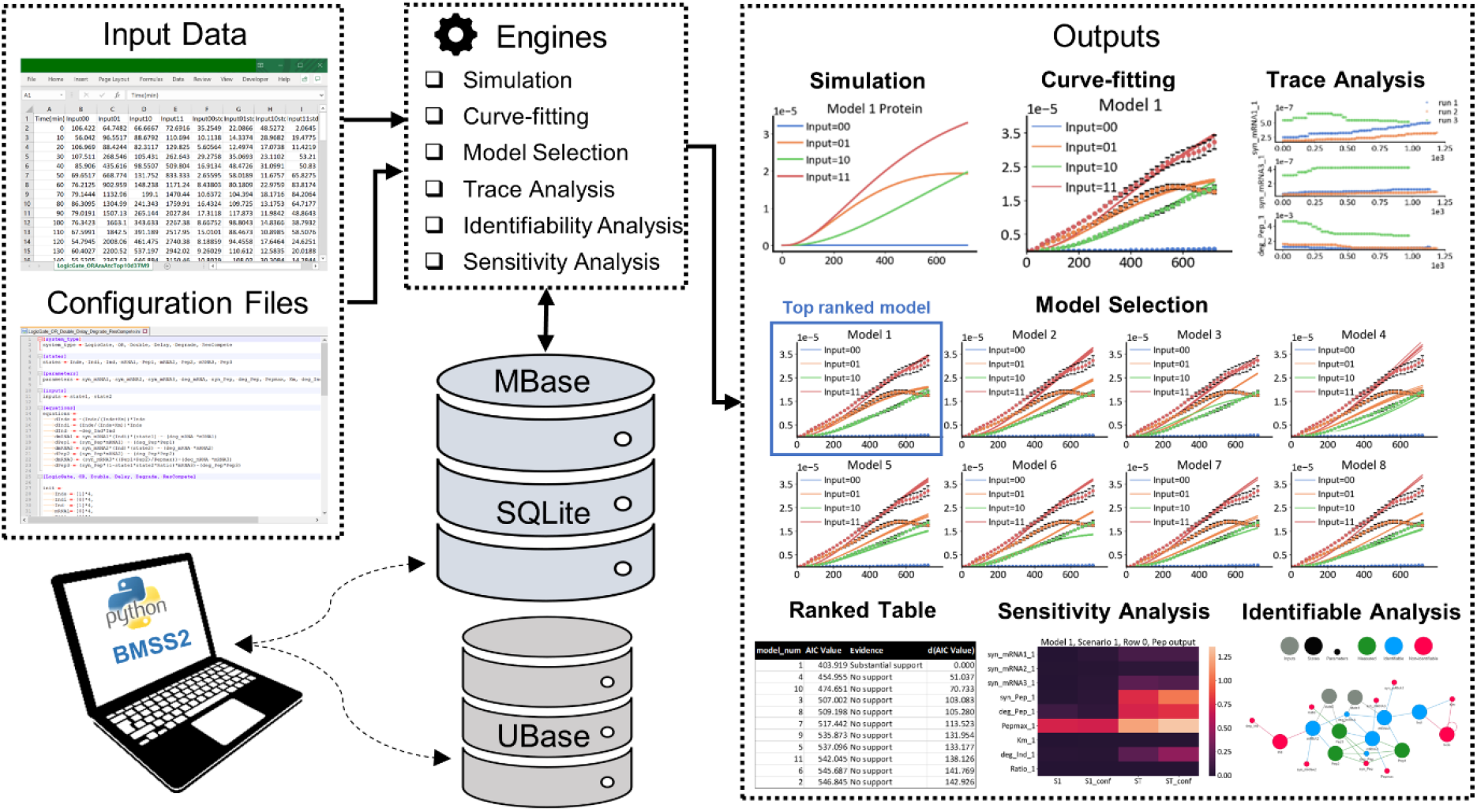
Overview of BMSS2: Models can be read from configuration files, retrieved from the global MBase or the local UBase. Operations can then be implemented on the models using modules from the BMSS2 core engine. Experimental input data can be imported for model fitting and model selection. The outputs exemplified an OR gate genetic circuit with one input driven by arabinose-pBAD promoter and the other by aTc-pTet promoter, while both were expressing red fluorescent proteins.

To model a biological system or gene device in forward or reverse engineering approach, users can either retrieve a pre-built model from MBase or UBase or formulate their own models using highly human-readable configuration files. *A priori* identifiability analysis can be performed on candidate models using our Python implementation of the STRIKE-GOLDD algorithm originally developed in MATLAB (Villaverde et al., 2019) to highlight unidentifiable parameters and unobservable states. Using this information, models can be reformulated, and experiments can be designed to acquire data to enhance parameters identifiability and model predictability. For parameter estimation, models can be fitted to experimental data using global optimizers (e.g., Bayesian inference) and practical identifiability can be verified using trace analysis. Finally, from a library of models, the users can apply the automated model selection feature to select the model that best balances goodness-of-fit and complexity using AIC calculation. With the top ranked model, SALib’s global sensitivity analysis (Herman and Usher, 2018) can be implemented to identify sensitive parameters for tuning based upon the desired outputs. Model prediction can then be executed to simulate the new system behaviours. The optimized model can ultimately be stored or updated iteratively in the local database for future reuse or contributed to the global MBase by submitting to the authors for community usage.

To demonstrate the utilities of BMSS2, we have applied the different functionalities spanning across various systems including inducible systems, logic gates, bacterial growth models and cellular resource model (Weiße et al., 2015).

## Acknowledgement

This work was supported by Singapore National Research Foundation [NRF SBP-P6 and SBP-P5], the Synthetic Biology Initiative of the National University of Singapore [DPRT/943/09/14], Summit Research Program of the National University Health System [NUHSRO/2016/053/SRP/05], and NUS startup grant [R-397-000-257-133].

## References

Aoki, S.K. et al. (2019). A universal biomolecular integral feedback controller for robust perfect adaptation. Nature, 570, 533–537.

Caliendo, F. et al. (2019). Engineered cell-based therapeutics: Synthetic biology meets immunology. Front. Bioeng. Biotechnol., 7, 1–8.

Choi, K. et al. (2018). Tellurium: An extensible python-based modeling environment for systems and synthetic biology. Biosystems, 171, 74–79.

Elowitz. (2000). A synthetic oscillatory network repressilator. Nature, 403, 335–338.

Herman, J. and Usher, W. (2018). SALib: An open-source Python library for Sensitivity Analysis. J. Open Source Softw., 41, 2015–2017.

Holowko, M.B. et al. (2016). Biosensing Vibrio cholerae with Genetically Engineered Escherichia coli. ACS Synth. Biol., 5, 1275–1283.

Hoops, S. et al. (2006). COPASI - A COmplex PAthway SImulator. Bioinformatics, 22, 3067–3074.

Jayaraman, P. et al. (2016). Blue light-mediated transcriptional activation and repression of gene expression in bacteria. Nucleic Acids Res., 44, 6994–7005.

Jayaraman, P. et al. (2018). Programming the Dynamic Control of Bacterial Gene Expression with a Chimeric Ligand- and Light-Based Promoter System. ACS Synth. Biol., 7, 2627–2639.

Myers, C.R. et al. (2007). Python unleashed on systems biology. Comput. Sci. Eng., 9, 34–37.

Olivier, B.G. and Snoep, J.L. (2004). Web-based kinetic modelling using JWS Online. Bioinformatics, 20, 2143–2144.

Villaverde, A.F. et al. (2019). Full observability and estimation of unknown inputs, states and parameters of nonlinear biological models. J. R. Soc. Interface, 16, 20190043.

Watanabe, L. et al. (2019). IBIOSIM 3: A Tool for Model-Based Genetic Circuit Design. ACS Synth. Biol., 8, 1560–1563.

Weiße, A.Y. et al. (2015). Mechanistic links between cellular trade-offs, gene expression, and growth. Proc. Natl. Acad. Sci. U. S. A., 112, E1038–E1047.

Yeoh, J.W. et al. (2020). A model-driven approach towards rational microbial bioprocess optimization. Biotechnol. Bioeng., 1–14.

Yeoh, J.W. et al. (2019). An Automated Biomodel Selection System (BMSS) for Gene Circuit Designs. ACS Synth. Biol., 8, 1484–1497.

