## Supplementary Data for "BMSS2: a unified database-driven modelling tool for systematic model selection and identifiability analysis"

### 1. Database

The BMSS2 database is implemented as two .db files; MBase, which stores curated models that come as part of the download; UBase which stores models created by the user for local use. Only UBase can be updated or modified via regular front-end function calls. Users can submit new models to us upon which we will add them to the latest version of MBase after curation. MBase is also available using our web application. This allows users to access the latest models added.

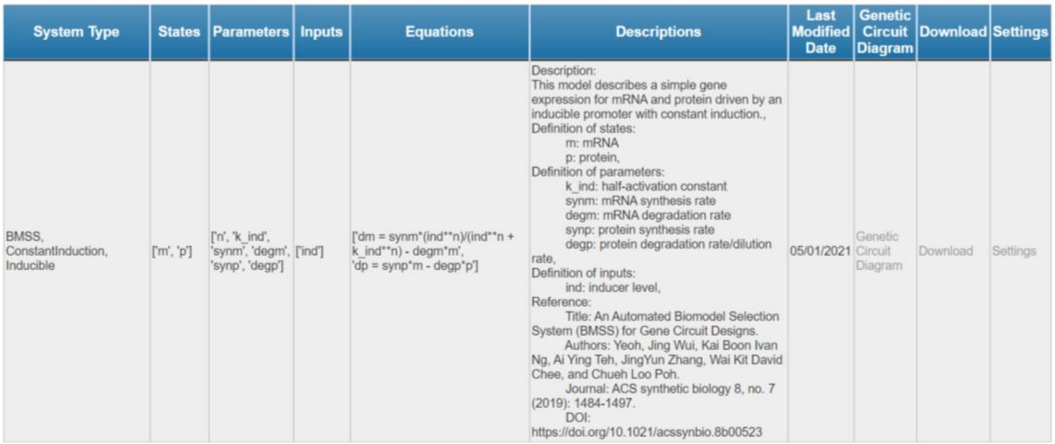

| System Type | States | Parameters | Inputs | Equations | Descriptions | Last Modified Date | Genetic Circuit Diagram | Download | Settings |
|---|---|---|---|---|---|---|---|---|---|
| BMSS, ConstantInduction, Inducible | ['m', 'p'] | ['n', 'k_ind', 'synm', 'degm', 'synp', 'degp'] | ['ind'] | ['dm = synm*(ind**n)/(ind**n + k_ind**n) - degm*m', 'dp = synp*m - degp*p'] | Description: This model describes a simple gene expression for mRNA and protein driven by an inducible promoter with constant induction., Definition of states: m: mRNA p: protein, Definition of parameters: k_ind: half-activation constant synm: mRNA synthesis rate degm: mRNA degradation rate synp: protein synthesis rate degp: protein degradation rate/dilution rate, Definition of inputs: ind: inducer level, Reference: Title: An Automated Biomodel Selection System (BMSS) for Gene Circuit Designs. Authors: Yeoh, Jing Wui, Kai Boon Ivan Ng, Ai Ying Teh, JingYun Zhang, Wai Kit David Chee, and Chueh Loo Poh. Journal: ACS synthetic biology 8, no. 7 (2019): 1484-1497. DOI: https://doi.org/10.1021/acssynbio.8b00523 | 05/01/2021 | Genetic Circuit Diagram | Download | Settings |

**Figure 1:** A screenshot of a model for an inducible system in the BMSS2 web application.

### 2. Configuration .ini Files

BMSS2 supports high-level functions that require different input arguments and data structures. While users can code these arguments themselves, this process can be tedious and prone to error. To simplify the process, BMSS2 provides a standardized solution by allowing users to use configuration .ini files to specify such information in a clear and streamlined manner. These information include (1) core models which represent the topology of a model; (2) settings for instantiating the arguments for simulation and analysis; and (3) template settings for storing default values for (2). For sample files, users may refer to the tutorials that come as part of the download.

### 2.1. Core Models

A core model data structure is a dictionary describing the topology of a model. The information for this data structure can be documented as shown below. In this example, a system called "MyModel, Version1" comprises two states, five parameters and one input.

```
[system_type]
system_type = MyModel, Version1

[states]
states = m, p

[parameters]
parameters = k_ind, synm, degm, synp, degp

[inputs]
inputs = ind

[equations]
equations =
        dm = synm*ind/(ind + k_ind) - degm*m
        dp = synp*m - degp*p
```

**Figure 2:** A core data structure specified in .ini format. The inputs section can be deleted if the model has no input variables.

### 2.2. Analysis Settings

**Simulation**: To simulate the time response of a model, users are required to specify the configurations for init (initial conditions), parameter_values (parameter values), and tspan (time span for simulation) for numerical integration in another .ini file or in the same file as the core model.

```
[MyModel, Version1]

init =
        m = [0],
        p = [0]

parameter_values =
        k_ind = [1e-2]*2,
        synm  = [1e-5]*2,
        degm  = [0.015]*2,
        synp  = [1e-2, 5e-2],
        degp  = [0.012]*2,
        ind   = [8e-2]*2

tspan =
        [0, 600, 31]
```

**Figure 3:** Simulation configurations specified in .ini format. Note that the input variable called "ind" has been combined with the parameters as BMSS treats inputs and parameters similarly during numerical integration.

Configurations can accommodate more than one set of conditions for numerical integration. In this example, the model is integrated with two values of synp.

```
[MyModel, Version1]

init =
        m = [0, 1e-5],
        p = [0, 1e-6]

parameter_values =
        k_ind = [1e-2]*2,
        synm  = [1e-5]*2,
        degm  = [0.015]*2,
        synp  = [1e-2, 5e-2],
        degp  = [0.012]*2,
        ind   = [8e-2]*2

tspan =
        [0, 600, 31]
```

**Figure 4:** Simulation configurations for integrating two sets of parameter values with two sets of initial conditions giving a total of four numerical integrations.

In this example, we want to consider two different sets of initial values of the model (known as "scenarios"). Each scenario is in turn integrated with two values of synp giving a total of four integrations. For more complex conditions such as step changes in inducer concentrations, BMSS2 also allows piecewise integration where the parameters and initial values can be modified before integrating each segment. However, the functions for this need to be coded separately.

**Sensitivity Analysis**: Users can test how perturbations in parameters affect the value of a particular objective function. The arguments for the configuration file are similar to those required for simulation but with a few extras. The fixed_parameters field allows one to specify parameters to be excluded from permutation. Note that the objective function needs to be defined separately in the user's Python code.

```
[MyModel, Version1]

init =
        m = [0]*3,
        p = [0]*3

parameter_values =
        k_ind = [1e-2],
        synm  = [1e-5],
        degm  = [0.015],
        synp  = [1e-2],
        degp  = [0.012],
        ind   = [8e-2]

parameter_bounds =
        k_ind = [1e-3, 1e-1],
        synm  = [1e-6, 1e-4],
        degm  = [1e-3, 1e-1]

fixed_parameters =
        [synp, degp, ind]

tspan =
        [0, 600, 31]
```

**Figure 5:** Configuration for sensitivity analysis is similar to those for simulation but extra information on parameter bounds and fixed parameters.

In this example, we want to perturb synm, degm and k_ind. Most of the fields are the same as those in simulation but we now need to specify which parameters are fixed.

**Curve-fitting and Parameter Estimation**: BMSS2 supports parameter estimation by fitting models against experimental data. The default algorithm is Bayesian inference simulated annealing. SciPy's differential_evolution, basinhopping and dual_annealing algorithms are also supported.

An initial guess is required for the parameters and inputs. Parameters with known values can be specified using the fixed_parameters field. Priors can be specified in the form of lower/upper bound pairs ("rectangular" priors) using the parameter_bounds keyword or as a mean/standard deviation pairs (Gaussian priors) using the priors keyword. Note that the .ini file does not include configurations for pre-processing the experimental data.

```
[MyModel, Version1]

init =
        m = [0]*3,
        p = [0]*3

guess =
        k_ind = [1e-2],
        synm  = [1e-5],
        degm  = [0.015],
        synp  = [1e-2],
        degp  = [0.012],
        ind   = [8e-2]

parameter_bounds =
        k_ind = [1e-3, 1e-1],
        synm  = [1e-6, 1e-4],
        degm  = [1e-3, 1e-1]

fixed_parameters =
        [synp, degp, ind]

tspan =
        [0, 600, 31]
```

**Figure 6:** Configurations for curve-fitting. In this example, we have three sets of data (i.e., three "scenarios"). The initial values are thus repeated thrice. The parameter_values section has been replaced with a guess section which will be used as the starting point for simulated annealing while the parameter_bounds section limits the parameter values for sampling. Finally, fixed_parameters tells BMSS which parameters are already known.

In this example, we want to estimate three parameters of the inducible system; k_ind, the Michaelis constant for induction; synm, the maximal mRNA synthesis rate; degm, the rate constant for mRNA degradation. We therefore list the remaining parameters and inputs under the fixed_parameters field while populating the parameter_bounds and priors fields to constrain the unknown parameters.

**A Priori Identifiability Analysis**: BMSS2 allows one to use python version of the STRIKE-GOLDD algorithm developed by Villaverde et al., originally in MATLAB to perform *a priori identifiability analysis* to highlight unidentifiable parameters and unmeasurable states. This could improve the model predictability and enable the users to better apply the model-driven approach to guide their experimental designs.

```
[MyModel, Version1]

init =
        m = 1,
        p = 1

input_conditions =
        Ind = 3

measured_states =
        [p]

parameter_values =
        synp = 1e-2,
        degp = 0.012
```

**Figure 7:** Configurations for running STRIKE-GOLDD. The input variable "Ind" should not be put under parameter_values as required by the STRIKE-GOLDD algorithm.

The value of the input_conditions field is equal to the number of inducer concentrations. If the inducer concentration changes continuously (such as a degrading inducer), the value should be "inf" without quotation marks. Meanwhile the measured_states field contains a list of states that are to be measured during the experiment. The parameter_values field contains values of known parameters while the init field contains the initial values of the states. Note that only one value for each state/parameter is allowed.

### 2.3. Template Settings

BMSS2 is able to store default values that can be used to generate template files for the analysis settings as shown below.

```
[settings1]

system_type = MyModel, Version1

init =
        m = [0],
        p = [0]

guess_ =
        k_ind = [1e-2],
        synm  = [1e-5],
        degm  = [0.015],
        synp  = [1e-2],
        degp  = [0.012],
        ind   = [8e-2]

fixed_parameters =
        [k_ind, synp, degp]

tspan =
        [0, 600, 61]

units =
        k_ind = molL-1,
        synm  = molL-1min-1,
        degm  = min-1,
        synp  = min-1,
        degp  = min-1,
        ind   = molL-1
```

**Figure 8:** BMSS can store default values of settings that can be used to generate templates. When creating a default, the corresponding core model is specified in the system_type subsection.

When creating settings templates, the section name is the name of the template. Meanwhile, the system_type is specified as a subsection.

### 3. SBML

BMSS2 can combine core model and default settings information to produce SBML files for interoperability with other software.

### 4. Model Selection

BMSS2 allows users to fit multiple candidate models to experimental data after which the best parameter sets of the multiple models can be used for model selection. BMSS2's functions for ranking models make use of the Akaike Information Criterion defined as

$$2n_{parameters} - 2\ln{(log\_likelihood)}$$

In the case of a negative log likelihood, the minus sign is moved into the brackets so that we obtain

$$2n_{parameters} + 2\ln{(-log\_likelihood)}$$
